# Lethal Haplotypes and Candidate Causal Mutations in Angus Cattle

**DOI:** 10.1101/125203

**Authors:** Jesse L. Hoff, Jared E. Decker, Robert D. Schnabel, Jeremy F. Taylor

## Abstract

**Background:** If unmanaged, high rates of inbreeding in livestock populations adversely impact their reproductive fitness. In beef cattle, historical selection strategies have increased the frequency of several segregating fatal autosomal recessive polymorphisms. Selective breeding has also decreased the extent of haplotypic diversity genome-wide. By identifying haplotypes for which homozygotes are not observed but would be expected based on their frequency, developmentally lethal recessive loci can be localized. This analysis comes without the need for observation of the loss-associated phenotype (e.g., failure to implant, first trimester abortion, deformity at birth). In this study, haplotypes were estimated for 3,961 registered Angus individuals using 52,545 SNP loci using findhap v2, which exploited the complex pedigree among the individuals in this population.

**Results:** Seven loci were detected to possess haplotypes that were not observed in homozygous form despite a sufficiently high frequency and pedigree-based expectation of homozygote occurrence. These haplotypes were identified as candidates for harboring autosomal recessive lethal alleles. Of the genotyped individuals, 109 were resequenced to an average 27X depth of coverage to identify putative loss-of-function alleles genome-wide and had variants called using a custom in-house developed pipeline. For the candidate lethal-harboring haplotypes present in these bulls, sequence-called genotypes were used to identify concordant variants. In addition, whole-genome sequence imputation of variants was performed into the set of 3,961 genotyped animals using the 109 resequenced animals to identify candidate lethal recessive variants at the seven loci.

**Conclusions:** Selective breeding programs could utilize the predicted lethal haplotypes associated with SNP genotypes. Sequencing and other methods for identifying the causal variants underlying these haplotypes can allow for more efficient methods of management such as gene editing. These two methods in total will reduce the negative impacts of inbreeding on fertility and maximize overall genetic gains.

## Background

The implementation of a national animal evaluation system in U.S. registered Angus cattle has generated estimates of genetic merit that are used to evaluate and select elite seedstock. Selection on individual traits and indices of traits has resulted in the genetic improvement of multiple traits, such as growth rate, carcass quality and calving ease [1, 2]. At the same time, artificial insemination has increased the utilization of certain paternal lineages. Selective breeding in livestock is known to contribute to the enrichment of deleterious alleles carried by highly utilized sires and also to increase the overall levels of relatedness among individuals. Many numerically important breeds, such as Holstein, Jersey, Nordic Red, and Angus, with extensive use of artificial insemination have recently found autosomal recessive lethal loci at moderate frequencies and that significantly impact fertility [3]. In recent decades several defects that are effectively lethal such as, Neuropathic hydrocephalus, Arthrogryposis Multiplex and Osteoperosis have been propagated within international Angus populations [4, 5]. A striking feature of these defects is their high prevalence in the U.S. registered Angus population, despite their severely deleterious phenotypic presentation [4]. This suggests that it is possible for recessive loci to have major negative impacts on fertility and overall performance without their early detection [3–5]. Consequently, recessive loci causing embryonic loss early in gestation could exist at relatively high population allele frequencies. These loci can reach high frequencies due to drift following severe population bottlenecks, due to their propagation by the extensive use of popular sire lines via artificial insemination, or due to linkage to beneficial alleles at strongly selected loci [7]. The extent of the impact that these alleles have on fertility and fitness in livestock is unknown, particularly because most of the reproductive process is unmonitored. However, when these impacts become large, they can be detected by genomic analysis of the population. In this study, we examined the inheritance of haplotypes genome-wide in U.S. registered Angus cattle to reveal candidate haplotypes that may harbor variants that cause embryonic lethality.

The dynamics of inbreeding depression in closed populations can be described statistically, but genomic analysis can reveal their biological basis. The accumulation and impacts of inbreeding are a function of the effective population size, the initial genetic load of deleterious sites, mating patterns, strength of selection and the extent or range of linkage disequilibrium within the genome [8]. Our work focuses on sites for which fitness = 0 when homozygous for a deleterious allele. Consequently, identity by descent at loci harboring loss of function (LOF) alleles can significantly impact fitness in populations that are accumulating inbreeding [9]. What is unclear, is how much inbreeding is tolerable and how common are recessive lethal alleles in livestock. Recent investigations in inbred human populations suggest that high levels of relatedness are needed to significantly impact fitness. Counterintuitively, up to a certain high threshold, parental relatedness appears to be beneficial for fitness [10]. An investigation into the North American human Hutterite population estimated that an average of 0.29 recessive lethal variants exist per haploid human genome, and that the primary force that removes these alleles from a population is drift [11]. Simulations by these authors also revealed that the majority (57.4%) of recessive deleterious variants that were segregating in the founder population were not observed in the modern population. Of the remaining recessive variants, only 8.23% had phenotypic effects. This result is important, because it suggests that in a population such as the U.S. registered Angus breed, a high proportion of latent recessive lethal variants could still be segregating without their having been detected by breeders.

A recent study in U.S. Holstein dairy cattle using genotyped trios found that expected inbreeding coefficients of offspring (obtained by simulated matings of actual parental genotypes) were slightly lower than the realized inbreeding coefficients, suggesting that increased inbreeding was not a constraint on viability of offspring [12]. This result, along with the knowledge of the nature of several existing recessive defects in the U.S. Holstein population suggests that the frequency of lethals may not be sufficiently high to produce an observable impact on the population when the average inbreeding coefficient is only 3.53%. These findings are also consistent with a recent study of dog domestication and breed formation that found that the major factor underlying the enrichment of deleterious variation in modern breed dogs was severe population bottlenecks, and not recent inbreeding [13].

Given experiences in other breeds and species, we hypothesize that there may be many more recessive lethal alleles in the U.S. registered Angus population than have previously been detected. To identify these, we implemented a method that was first described by VanRaden [14]. A sample of 3,961 genotyped Angus cattle, primarily bulls extensively used in artificial insemination that were members of a pedigree containing 117,212 identified individuals, was analyzed to identify haplotypes that were expected to be observed but that were not actually observed in the homozygous state. The pedigree spanned more than 60 generations with the earliest ancestor born in 1836, and with the earliest genotyped animal born in 1955 [2]. The analysis assumed that the haplotypes harbored fully penetrant lethal alleles that would preclude the viability and genotyping of an animal homozygous for the haplotype. Within this pedigree, haplotypes that were identical by descent were repeatedly sampled, allowing for the detection of autosomal recessive lethal alleles.

The adoption of high-density array-based genotyping in commercial beef and dairy cattle populations is rapidly increasing, due to the utility of genomic selection [15] and as a consequence a large number of trios, patrios (sire, maternal grandsire and son) and half-sib families have now been genotyped. Whole-genome sequencing of influential population members and the use of genotype imputation will allow the identification of lethal alleles without the observation of the phenotype responsible for the loss [16]. Characterizing the number and identity of these variants will provide a deeper understanding of the biological and quantitative underpinnings of inbreeding depression. It will also enable enhanced management of animal reproduction, as these variants can be identified in any genotyped animal.

## Methods

### Genotypes and Animals

BovineSNP50 BeadChip (Illumina, San Diego, CA)[17] data for 3,993 registered Angus animals born between 1955 and 2012 and representing 63 generations were available for analysis. Genotypes had been filtered to retain data with a call rate of ≥ 90% and minor allele frequency (MAF) ≥ 0.01 [2]. Pedigrees had also been validated by homozygous transmission incompatibility rates estimated as the frequency of loci for which the parent and offspring were alternate homozygotes. Pedigree relationships were expunged when the incompatibility rate exceeded 1.5%, and individuals who did not match their recorded parents or offspring were removed from the pedigree. After filtering, 52,545 loci and 3,961 animals remained. These sites were next phased and missing genotypes were imputed using findhap (v2) based upon a combination of simple haplotype frequency sorting and the use of all available pedigree information [18].

### Population and Pedigree Based Haplotype Analyses

Haplotypes were defined by 20-marker sliding windows genome-wide. Evidence for a lethal allele within a haplotype was evaluated using two statistics. The first was based on population frequency. For each haplotype in each window, frequency was estimated and the number of individuals in the population that were homozygous for the haplotype was tallied. When there was an absence of homozygotes, the likelihood of this occurrence was calculated under a model that assumed carriers to be randomly mating and selective neutrality of the haplotype.

In the second analysis, the actual matings within the pedigree were used to estimate the expected number of homozygotes for any given haplotype based upon the number of matings involving carriers and the assumption of selective neutrality. The pedigree for the genotyped animals was parsed to identify patrios, defined here as families for which genotypes were available for the offspring, sire and maternal grandsire. Maternal genotypes were rarely available as DNA was primarily sourced from cryopreserved semen [2]. This family structure allowed the testing of segregation distortion when both the sire and maternal grandsire were heterozygous for the same haplotype. For each sliding window of 20 markers if a haplotype was never observed in the homozygous state and had a sample frequency of greater than 2%, the number of families for which the sire and maternal grandsire were both heterozygous was counted. The probability of observing at least one homozygote was then calculated based on the count of patrios and the assumption of selective neutrality. The probability that no homozygous *hh* haplotypes are observed in the progeny of C patrios when both the sire and maternal grand-sire are *Hh* heterozygotes and the *h* haplotype does not harbor a selected deleterious allele is (0.875 - 0.25q)^c^ where q is the frequency of the *h* haplotype in the population. There were 2,480 patrios represented in the sample. Regions of the genome with an identified deficit of homozygotes for a specific haplotype were examined for underlying genes using pybedtools and the UMD3.1 genome assembly annotation run 104 [14, 15].

### Generation of Sequence Data

To further analyze the variation within the genomic regions harboring haplotypes that were deficient for homozygotes, the whole-genome sequences of 109 animals from this population were examined [20]. These animals were selected for sequencing based upon their impact on the breed assessed by the expected numbers of genome equivalents (progeny have 0.5, grandprogeny have 0.25, etc) present within registered descendants in the population. They were sequenced with paired-end 2 × 100 bp sequence reads to an average of 27X depth of coverage of the UMD3.1 assembly with Illumina Genome Analyzer, GAII, HiSeq 2000 or 2500 instruments from two libraries with 350 bp and 550 bp average fragment sizes. FastQC was used to analyze the quality of the reads [21]. Exact duplicates were removed, and adapters were trimmed using a custom in-house Perl script. All remaining reads were error corrected using QuorUM [22]. Newly created duplicates (due to the trimming of low quality ends and correction of errors) and reads shorter than 35 bp were removed and the final data set was aligned to the UMD3.1 reference genome assembly using NextGENe 2.4.1 (SoftGenetics, LLC, State College, PA) alignment software. Reads were required to have a matching segment at least 35 bases long and 95% overall match, and a maximum of 2 bases of mismatch across the whole alignment. Up to 1000 alignments of equal likelihood were allowed genome-wide. NextGene 2.4.1 was also used for variant calling.

The sequence-derived variants for the 109 bulls were used for two purposes; identification of variants shared amongst animals identified as carriers for the putatively lethal BovineSNP50 20 SNP haplotypes, and for the imputation of the entire genotyped population to whole-genome sequence variation.

### Examining Carrier Sequence Data

Within the genomic regions identified from the marker data as containing candidate lethal haplotypes, all variants identified in our sequenced sample were analyzed for carrier concordance, implemented using python scripts. This involved examination of variants that were observed as heterozygous in all bulls predicted to be carriers of the homozygote deficient haplotype. The sample size for the sequenced animals was not sufficient to run the population allele frequency or patrio analyses.

### Imputation of BovineSNP50 Genotypes to Whole Genome Sequence Variation

We imputed the BovineSNP50 genotypes for this population to whole-genome sequence level variation in order to identify potential lethal variants, using the 109 sequenced animals as the reference population. We selected 24,974,785 SNPs from the full set that had been identified in these animals. We first included SNPs found in the 109 sequenced Angus bulls that were biallelic and located within gene boundaries (777,432), UTRs (33,379) or that were splice site variants (636,492). These variants spanned the allele frequency spectrum but were identified at high sequence coverage. To enable imputation we also included 23,527,482 variants that had been independently identified in run 5 of the 1000 Bull Genomes project [23]. These variants represent filtered, high quality segregating sites identified from the whole genome sequences of 1,578 animals from multiple taurine breeds including Angus. Genotypes called for the 24,974,785 variable sites genome-wide in the 109 registered Angus bulls were used as the reference set for whole-genome sequence imputation of the BovineSNP50 data using Fimpute [24].

The imputed genotypes were individually analyzed for the absence of homozygotes for alleles present within each of the candidate haplotyped loci using the frequency and pedigree approaches. We first identified high frequency variants for which no homozygous individuals were predicted. The pedigree analysis was also performed for all candidate variants identified in the frequency analysis. Variants identified by either of these processes were characterized using the variant effect predictor release 79 [25].

## Results

### Identification of Putative Lethal Haplotypes

We identified seven haplotypes with a pattern of inheritance in U.S. registered Angus cattle that suggests that they each harbor an autosomal recessive lethal allele (Table 1). Using a binomial distribution for the number of observed homozygotes in the progeny of C patrios, we calculated the probability of observing no homozygotes when each haplotype was selectively neutral. We selected haplotypes as putatively harboring autosomal recessive lethals when the probability of observing no homozygotes was less than 0.02. This threshold for statistical significance provides considerable confidence that the lack of homozygosity for these haplotypes did not occur by chance alone.

**Table 1:** Chromosomal regions predicted to harbor lethal haplotypes identified in the analysis of the BovineSNP50 data.

| Chr | Location (Mb) | Length (Mb) | Haplotype Frequency<sup>1</sup> | Number of Patrios<sup>2</sup> | Probability<sup>3</sup> | Sequenced Carriers | Concordant Variants | Concordant In High Coverage |
| --- | --- | --- | --- | --- | --- | --- | --- | --- |
| 1 | 27.7-29.0 | 1.3 | 0.023 | 39 | 0.0042 | 1 | 4 | 4 |
| 4 | 82.5-84.0 | 1.5 | 0.076 | 127 | 2.66E-09 | 21 | 9 | 118 |
| 8 | 62.0-63.0 | 1.0 | 0.023 | 35 | 0.0074 | 5 | 1 | 1 |
| 12 | 60.0-61.2 | 1.2 | 0.032 | 46 | 0.0014 | 12 | 0 | 0 |
| 15 | 82.3-83.1 | 0.8 | 0.038 | 31 | 0.011 | 10 | 1 | 1 |
| 17 | 46.5-47.5 | 1.0 | 0.045 | 49 | 0.00076 | 15 | 2 | 2 |
| 29 | 43.0-44.2 | 1.2 | 0.044 | 118 | 3.22E-08 | 16 | 3 | 13 |

<sup>1</sup>Haplotypes estimated for 20 contiguous SNP loci. <sup>2</sup>Number of families out of 2,480 for which the sire and maternal grandsire were both heterozygotes for the haplotype. <sup>3</sup>Probability of observing no homozygous progeny if the haplotype is selectively neutral.

### Sensitivity to Window Size

We evaluated the sensitivity of identification of these marker-based haplotypes to window size and concluded that a window size of 20 contiguous BovineSNP50 markers was appropriate for capturing the haplotypic diversity within the population (Figure 1). This window size appears to discriminate between haplotypes that are identical by descent (IBD) and those that are identical by state (IBS). Our analysis shows that the number of common haplotypes detected rises as window size increases, but begins to plateau at 20 markers. Considering the moderate marker density of the BovineSNP50 (1 SNP per 50 kb), haplotypes that are defined by only two markers are assumed to be IBS for the purposes of analysis but likely actually represent a number of distinct haplotypes at the level of genome sequence. As the window size increases, the likelihood increases that two haplotypes found in different individuals that are IBS are also IBD and are thus concordant at the level of sequence variation. However, with large window sizes, recombination may lead to a lethal variant being present on more than one haplotype, thus decreasing the power of the analysis. Indeed, as window size increases, the overall rate of individuals homozygous for any haplotype declined. The window size selected for this study appears to achieve an appropriate balance of genome-wide homozygosity and rate of occurrence of high frequency haplotypes (Figure 1). Slight changes in the haplotype window size did not affect the detection of the 7 haplotypes reported in Table 1.

**Figure 1:**
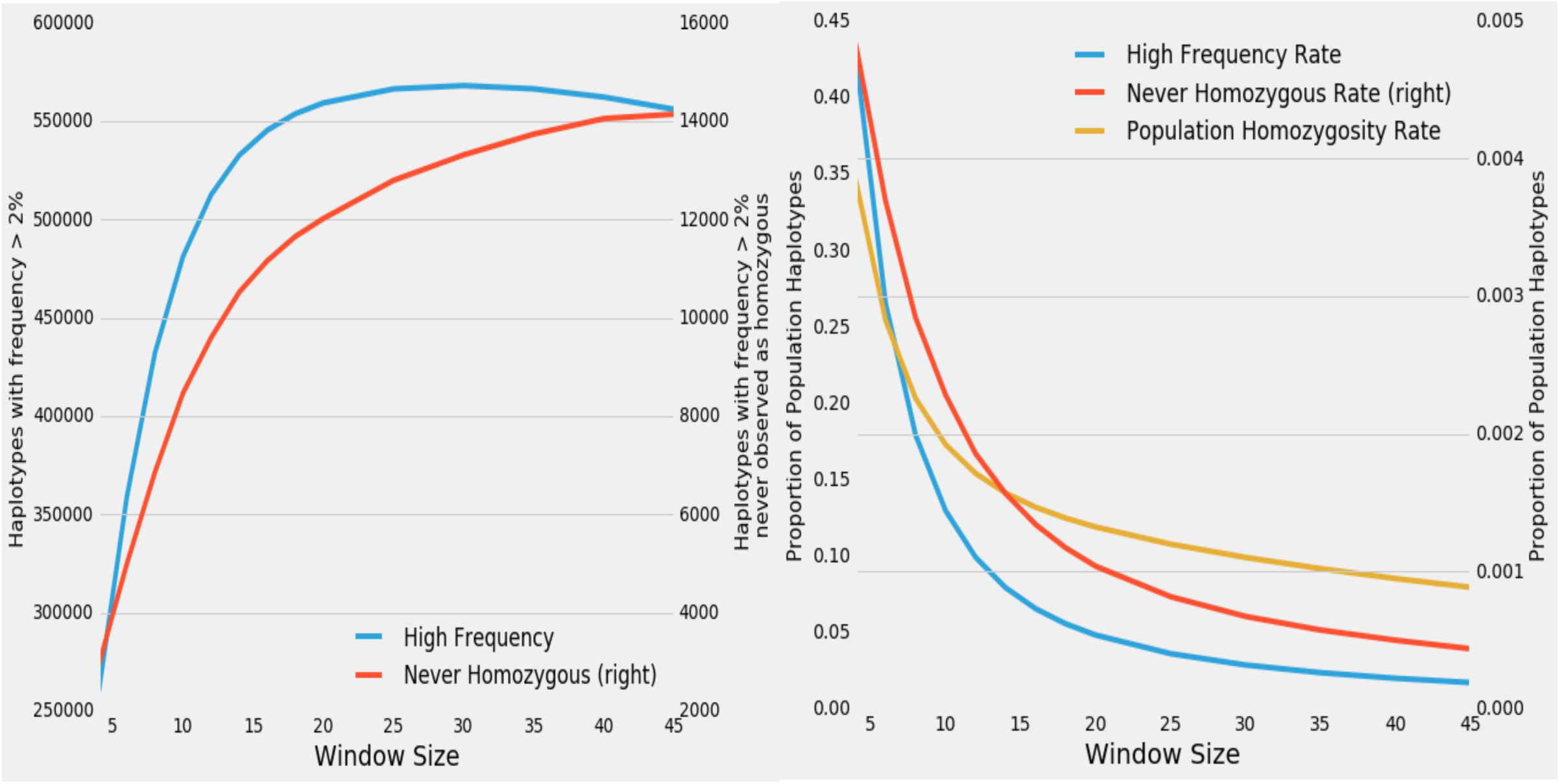
Effect of window size on haplotypic diversity and lethal haplotype detection. A) As the size of the window expands, many more distinct haplotypes are detected genome-wide. However, fewer of the newly detected haplotypes are common as window size increases, and the number of common haplotypes that are never observed as being homozygous asymptotes. B) Rate of homozygosity, which is the percentage of individuals that are homozygous for any haplotype, is high for small window sizes but quickly declines. The assumption that phased marker homozygosity implies identity by descent underlies the population frequency and patrio tests for haplotype lethality.

We were also able to validate IBD status of these regions by an examination of the sequence data generated for animals that were predicted to be carriers of identical haplotypes. For each of the loci predicted to harbor recessive lethal haplotypes (Table 1), we identified the bulls among the 109 sequenced animals that were predicted to be carriers of each putative lethal haplotype and computed the pairwise rate of opposing homozygous sequenced sites between all pairs of carrier animals. For example, for the haplotype on chromosome 29, with 16 predicted carriers we made 120 pairwise comparisons using all sequence called variant genotypes within the haplotype coordinates (Figure 2). Amongst individuals that share a common haplotype, the alternate homozygote rate is related to the error rate for sequence-based genotype calls for heterozygotes. The rate observed in our predicted carriers within our 6 testable haplotype regions was similar to the rate for sire-son pairs. One region (Chr 1) had only one sequenced predicted carrier and could not be evaluated. The majority of the carrier pair comparisons with rates of opposing homozygous sequenced sites >0.01 were caused by the inclusion in the analysis of 3 individuals that had been sequenced to averages depths of < 10X. Within the haplotype, opposing homozygous sequenced site rates were markedly lower for predicted carriers than for randomly selected individuals. When predicted carriers for a lethal haplotype were analyzed for a randomly selected 20 marker region elsewhere in the genome, they had opposing homozygote rates that were similar to those of the unrelated sire pairs. This indicates that the haplotypes generated from the BovineSNP50 data successfully identified genomic regions that were IBD at the level of the genome sequence. We also observed that this method is sensitive to the depth of sequence coverage due to the inaccurate identification of heterozygous loci as being homozygous when the alternate allele was never sequenced. In total, 16 animals had depths of sequence coverage of < 10X and most of the remaining animals (70) had > 20X. For a Sire-Son pair, low sequence coverage for at least one member of the pair led to a rate of homozygous inconsistency that was increased by an order of magnitude.

**Figure 2:**
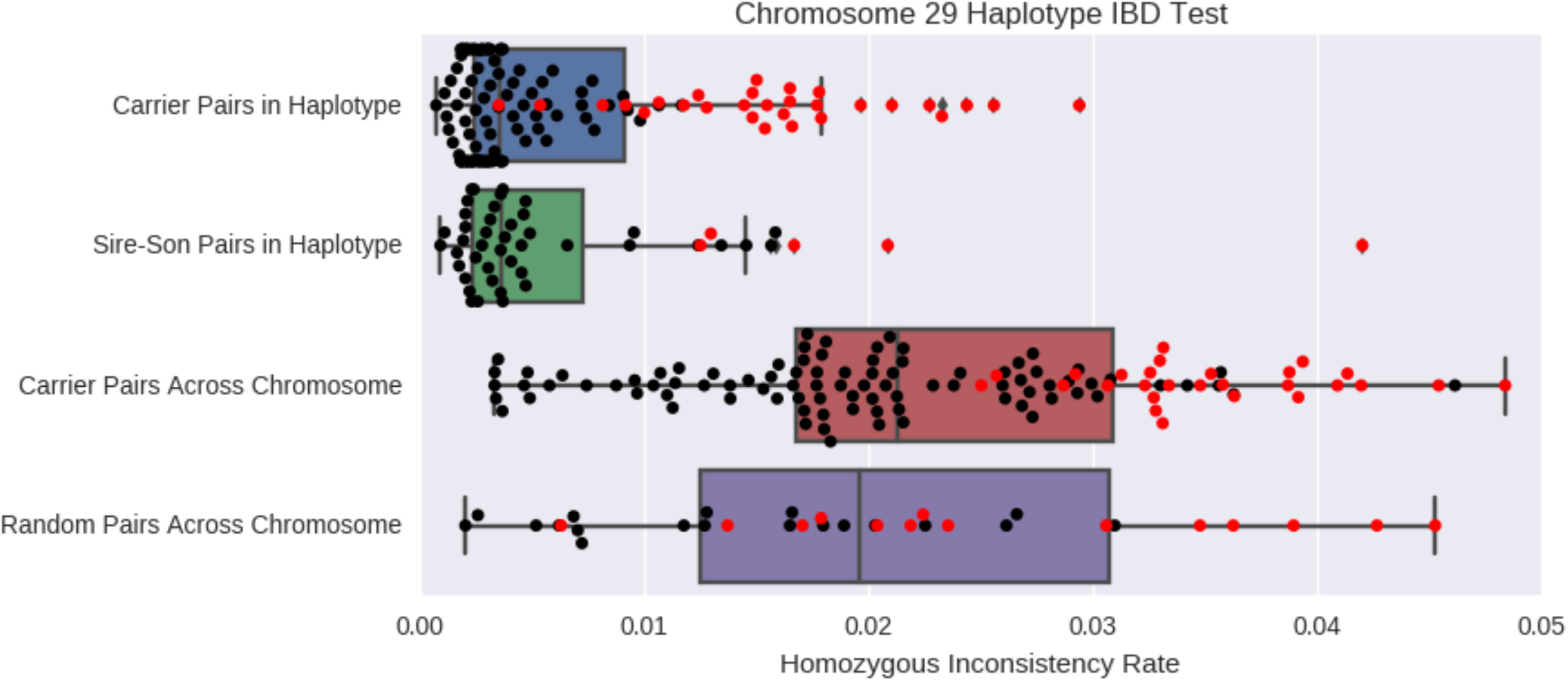
Validating the sequence level IBD status of bulls predicted to be carriers of a BovineSNP50 lethal haplotype using sequence data. Homozygous inconsistency rates are calculated pairwise amongst predicted carriers, sire-son pairs and randomly sampled animals (not 1^st^ degree relatives). Two different regions are compared: within the tested chromosome 29 haplotype and across the entirety of chromosome 29. The discordance rate among of pairs (shown in red) that contain one animal with low coverage (<10x) is greatly elevated.

### Analysis of Sequence Variation and Candidate Genes in Bulls Predicted to be Lethal Haplotype Carriers

To identify the causal variants underlying these putatively lethal haplotypes, we first directly examined resequencing data from bulls that were predicted to be carriers of the lethal haplotypes. Among the 109 sequenced bulls up to 21 animals were predicted to be carriers of each of the BovineSNP50 putatively lethal haplotypes reported in Table 1. Within these seven genomic windows, we identified candidate variants that were never homozygous in the 109 sequenced bulls. However, none were observed to be exclusively heterozygous in the predicted carrier animals. That is, all of the alleles found to be heterozygous in all of the predicted carriers were also observed to be heterozygous in animals that did not carry the predicted lethal haplotype. A recessive lethal variant could be heterozygous in animals that were not carriers of the putatively lethal haplotype if the mutation is sufficiently old that recombination has occurred relative to the haplotype on which the mutation occurred. Table 1 reports, for each genomic region predicted to harbor a lethal haplotype, the number of variants that were heterozygous in all predicted haplotype carriers and that were never found to be homozygous in any of the 109 resequenced bulls. For instance, on chromosome 15, an intronic variant in *SSRP1* was found in 9 of the 10 sequenced bulls that were predicted to carry the putatively lethal haplotype, and in 25 of all 109 sequenced Angus bulls, but was never found to be homozygous. Although the variant is intronic, with no expected impact on splice site variation, it resides in an interesting candidate gene as SSRP1 is essential for mouse embryonic development [26].

This analysis was also conducted after excluding the 16 animals sequenced to low average sequence depth of coverage (< 10X). Genotypes for these animals are likely to contain false positive homozygotes at many true heterozygous sites. This resulted in the detection of additional concordant loci but again none were exclusive to the predicted carriers. The chromosome 29 locus had 12 additional variants identified that were located in genes such as *CAPN, NRXN2, PACS1* and *PP25B*. However, none of these mutations are in exons or had other interpretations in Variant Effect Predictor (VEP) and all 12 were heterozygous in 18 animals, including 4 that were not predicted by their marker data to be carriers. Thus, the region appears to comprise a large consistent haplotype with no one particular variant being simply implicated as causal for lethality. The locus on chromosome 4 had 118 variants identified as being heterozygous in haplotype carriers following the removal of the low sequence coverage animals, but none were predicted to alter protein amino acid sequences.

The locus on chromosome 1 for which we predicted a lethal haplotype contains only one gene, *GBE1* that encodes a protein that catalyzes the branching of glycogen, the main form of energy storage in the body [24]. There was only one sequenced animal in our population predicted to carry this disease and further analysis of the unique heterozygotes within this gene were therefore not feasible.

### Analysis of Imputed Sequence Variation in 3,961 Registered Angus

When analyzed in the 3,961 animals, 2,504 of the 24,974,785 imputed variants had a MAF ≥ 2%, no predicted homozygotes, and were found in ≥ 30 double-carrier patrios. Of the 24,974,785 variants, 147,764 had a deleterious consequence predicted by their SIFT scores [30], but only 4 were among the 2,504 candidate autosomal lethal alleles. Two of these sites were in *LOC521645*, an olfactory receptor gene (near, but not within the chromosome 15 locus, Table 1), and another was in *LOC100336589* on chromosome 18. The fourth was on chromosome 21 in *GEMIN2*, for which a recessive lethal embryonic phenotype occurs in mouse knockouts and is associated with the survival motor neuron complex [31]. This allele had a frequency of 7.8% following imputation into the 3,961 animals and was observed in 55 double-carrier patrios without producing a homozygous progeny.

Twelve variants within *GBE1*, none of which were predicted to be deleterious, were at a MAF ≥ 2%, had no homozygotes predicted, and were found in ≥ 30 double-carrier patrios, with one variant present in 133 double-carrier patrios. However, only one of the sequenced animals was predicted to carry this haplotype making it difficult to use the data for the sequenced animals to exclude any heterozygous sites within this animal’s diplotype from candidacy for lethality. The haplotype on chromosome 1 spanning *GBE1* was the only predicted lethal to contain any of the 2,504 candidate variants identified in the analysis of the imputed whole genome sequence data.

## Discussion

Due to the repeated sampling of parental gametes and the sharing of haplotypes across families in this extensive Angus pedigree, we were able to powerfully test the fitness consequence of many haplotypes. The use of artificial insemination increases selection intensity by allowing relatively few individuals to sire large numbers of progeny and this population has historically been strongly selected for growth and calving ease [2]. Four bulls in this dataset were each represented in over 100 of the 2,480 genotyped patrios, suggesting that intense selection can drive deleterious alleles carried by these bulls to a relatively high frequency. In some cattle populations, allele frequencies of up to 20% have been observed for some recessive lethal haplotypes [6]. These common recessive lethals have the greatest impact on population fitness, but as genotyping becomes more pervasive, rare recessive lethal haplotypes will be detected and the available genotypes should allow for their management.

### Management Strategies

Angus breeders have historically selected against recessive lethal alleles which manifest as fatal calf defects in this population [4]. Registered animals are tested for known defects and registration is prevented for young animals that test to be carriers. As new deleterious alleles including those causing early embryonic loss are discovered, this approach is likely to be untenable. The seven new putatively autosomal recessive lethal haplotypes may now be predicted in hundreds of thousands of genotyped animals [32] and many more deleterious loci may be discovered as the number of genotyped animals increases and new lineages rise to prominence that contain yet to be detected recessive alleles. Management must shift from registration exclusion to a means of incorporating marker diagnostics into genomic selection and mate selection protocols.

A mate selection procedure has been implemented in the MateSel software that applies a linear weight against either the number of recessive alleles present in the progeny generation (LethalA) or the number of homozygous progeny (LethalG) [33]. In a simulation with a higher count (N=100) for the number of lethal alleles segregating in a population than identified in this study, it was not possible to sufficiently weight in either selection scheme to eliminate embryonic mortality. This scenario may approach reality as the number of identified deleterious recessive alleles causing calf defects as well as embryonic loss increases. These simulations also suggest that implementing the LethalG strategy is a more efficient means of achieving genetic gain while reducing the frequency of recessive alleles.

Cole (2015) suggested an alternative strategy in which parent average breeding indexes are adjusted for lost progeny by adding a cost associated with recessive haplotypes [34]. This method could be extended to account for the joint impact of multiple linked or unlinked segregating loci to reduce the frequency of deleterious alleles. Counterintuitively, this study also found a zero to inverse relationship between the embryo’s realized inbreeding coefficient and its probability of being homozygous for an allele responsible for a recessive disorder. This suggests that the goal of reducing the long-term rate of accumulation of inbreeding in breeding programs may not impact the rate of embryonic loss due to the action of recessive lethal alleles. This is consistent with the observation from studies of embryonic loss and coancestry in humans [8].

### Managing False Positives

Managing selection based on these results requires certainty about the lethal haplotype’s effect. In this study, a total of 12,020 haplotypes were found genome-wide (including partially overlapping haplotypes) that occurred at a MAF ≥ 2% but were never found as homozygotes. Assuming random mating, a haplotype at this frequency is expected to have, on average, only 1.5 homozygotes in our Angus sample. However, the existence of non-random mating within this population could substantially decrease the likelihood of observing homozygotes, and explain why so many of these regions were observed. The patrio analysis directly incorporates the matings that created the population to detect deviations from selective neutrality in progeny genotypes. However, this approach is limited by the availability of genotyped patrios. In breeds where insemination and pregnancy records are more detailed, these have been crucial for validating putative lethal haplotypes[14].

In the absence of these data, both the MateSel approach and the index adjustment could be adapted to account for the uncertainty of the lethality of the predicted allele or haplotype. In addition to validated haplotypes, and the high confidence haplotypes that we observe here, this approach could enable the incorporation of haplotypes into the selection scheme that appear to be lethal but that are at low frequency in the population. There are now Angus pedigrees in which hundreds of thousands of animals have been genotyped world-wide and an analysis of these data would likely improve the resolution of lethal haplotype detection and could also identify many rarer variants [15]. If these lethal alleles are individually rare but each individual carries many of them, recessive lethals could affect a substantial portion of pregnancies.

### False Negatives

Even with adequate sample sizes to ensure statistical power, there are limitations to the methods that we have employed. The approaches employed will only detect recessive alleles that are perfectly concordant with a BovineSNP50 haplotype. If a recent autosomal recessive lethal mutation has occurred on a common haplotype, the population will comprise haplotypes harboring either the lethal mutation or the wild type allele and homozygotes that include the wild type allele will be observed. In this case, very large sample sizes are required to detect deviations from the number of homozygotes expected under Hardy-Weinberg equilibrium. This could also involve restricting the analysis to different pedigree lineages for an IBS haplotype. This may explain why we failed to identify any of the recent recessive genetic defects found in the Angus breed [5]. In recent years, the alleles responsible for these defects have frequencies that have ranged from 3-9% [4]. However, the regions of the genome that harbor these alleles were not detected in the marker-based haplotype analysis and the causal variants were not detected in the analysis of the imputed sequence data. The likely cause of this is that the haplotypes we examined did not always indicate the presence of the defective allele. For instance, the mutation causing Neuropathic Hydrocephalus, originated in bull G A R Precision 1680 born in 1990 [35]. Consequently, there are both wild-type and deleterious versions of the BovineSNP50 haplotype on which the mutation arose segregating in the genotyped population. When larger sample sizes become available, it would be useful to repeat these analyses and test for homozygote deficiency rather than complete absence.

We have also not attempted to model mutations in loci with parent of origin affects, such as imprinting associated defects [36]. SNP array genotypes identify large heterozygous deletions as being homozygous for the alleles present on the non-deletion chromosome, which may prevent the identification of carriers for the deletion. This type of mutation has previously been associated with lethal recessive diseases in cattle [7]. Additionally, lethal alleles with incomplete penetrance will not be captured by our analyses. Other errors in genotyping, phasing or imputation also likely contribute to reductions in power to detect lethal haplotypes.

### Using sequence data

One potential solution to the limitations of array genotype data is to analyze sequence-derived and/or imputed genotypes. These data may help with the management of putative recessive lethal alleles. True causal variants can be tracked more effectively than marker haplotypes. When the causative alleles have been identified, gene editing may also present an efficient means of reducing the genetic load of elite sires in a manner that is complementary to the current breeding system [37].

Sequence data also has potential advantages for the detection of recessive loci. If appropriately processed to capture SNPs, large and small indels and structural variants, they directly represent the pool of all recessive deleterious alleles. However, in practice, sample sizes have been small and the identification of large indels, particularly insertions, and complex structural variants has been challenging. Our analysis of sequence data failed to identify any candidate causative mutations in the marker-based haplotypes that were predicted to be lethal. The sample size for the sequenced animals was not sufficient to conduct the frequency or pedigree analysis with the genome-wide sequence variants. Furthermore, we did not attempt to analyze many types of complex variation, such as large indels or structural variants [38]. Large structural variants are enriched for deleterious variation but can be complex to analyze with short read data [39]. Alternative analyses of the sequence data to identify these variants or the use of methods which generate longer reads may be necessary to capture the causative variants.

In this study we did not detect a particular variant within a putative haplotype that was likely to cause a recessive lethal phenotype. However, the gene within the region on Chromosome 1, *GBE1* appears quite promising. Mutations in this gene produce recessive phenotypes in mammals including horse, mouse and human [24, 25]. In the U.S. Quarter Horse population, phenotypes created by homozygotes for *GBE1* mutations ranged from stillbirth to early failure to thrive, with death never occurring later than 18 weeks of age [28]. Mouse knockout analysis revealed few visible or biochemical phenotypic effects in heterozygotes. Monitoring of embryonic development in homozygous knockouts revealed that deformities only occurred late in gestation and led to stillbirth or death shortly after birth. Mice with a construct with low GBE1 activity incorporated into their genome to replace the wild type allele demonstrated poor metabolic performance, and the accumulation of polyglucosan [29]. None of the mice with limited GBE1 function lived beyond 39 weeks, while all control mice survived the trial. While not all of the recessive *GBE1* genotypes in other species have resulted in embryonic loss, the reduced growth associated with homozygosity for these mutations makes it unlikely that an affected animal would be selected as a sire or dam. They would therefore be highly unlikely to be included in our genotyped sample of Angus cattle. However, identifying homozygous calves from those produced by mating carriers and assaying their GBE1 functionality might be possible.

### Mapping Candidate Variants without Sequencing

Rather than generating expensive sequence data for identifying recessive lethals, two strategies might be useful: assay development and imputation. Novel variants of all classes that are detected by sequencing can readily be incorporated onto commercial genotyping platforms. This expedites fine-mapping within a known lethal haplotype in a commercial population. Variants that had predicted deleterious functional impacts based on bioinformatic analysis would also be excellent candidates for inclusion on commercial genotyping platforms.

Imputation accuracy, particularly for rare variants, may not be sufficient to identify candidate segregating recessive lethals. We previously analyzed the accuracy of imputation using variants from Run4 of the 1000 bull genomes project as a reference for our Angus BovineSNP50 genotyped population using the same imputation methods that were used in this study [20]. Our 109 sequenced animals had their BovineSNP50 genotypes imputed to the 1000 Bull Genomes Run4 sequence reference set as well as variants called directly from their whole-genome sequences. Comparing these two sets of genotypes revealed correlations between genotypes in the range of 80-90% for common variants, and 60-80% for variants with MAF ≤ 10% [20]. We would expect the causal variants underlying these lethal haplotypes to fall in the rare, more inaccurately imputed frequency class. This greatly complicates the utility of imputation for this application, and the results of imputation presented here are not appropriate for application to breeding decisions. Had an interesting candidate locus with a plausible biological mechanism emerged, it would have been a good target for further confirmation and possibly immediate use. More sophisticated sequence imputation methods that provide higher imputation accuracies across the allele frequency spectrum are now becoming available. These have been used in human studies to identify individuals that are homozygous for rare variants with predicted deleterious effects [16]. Applying these methods to livestock populations with extensively described and genotyped pedigrees is feasible, and may prove useful for fine-mapping within candidate haplotypes.

## Conclusions

We identified 7 potentially recessive lethal haplotypes segregating in the U.S. registered Angus population that open opportunities for improving breeding success and increasing the mean fitness of the population. These haplotypes have been propagated throughout Angus lineages represented in a set of 3,961 genotyped animals and were not observed in homozygous form. The phenotypic effects of these haplotypes have not been directly observed, but may be inconspicuous such as in the event of early embryonic loss or may be unreported defects leading to the loss of the calf. Efforts to identify causal mutations with a clear molecular impact from sequence data were unsuccessful but interesting candidate genes such as *GBE1* were identified. Further validation of the impact of these haplotypes on fertility and the direct observation of calves that are homozygous for these haplotypes could reveal interesting biology. Our capacity to detect these loci will continue to improve as increasingly large numbers of animals are genotyped and sequenced. The quality of the reference genome assembly and methods for characterizing and imputing structural variants are also improving and will improve the quality of this type of analysis. Eventually, we will identify dozens of deleterious recessive loci, and can use chip-based genotyping to manage matings, track alleles through lineages and potentially use gene editing to remove them from elite animals.

## Declarations

### Ethics approval and consent to participate

The genotype data described in this manuscript was previously analyzed or collected from commercially generated animal semen; as such no ethics or animal welfare approval was required.

### Consent for publication

Not applicable

### Availability of data and material

Genotypes are available to scientists interested in non-commercial research upon signing a Materials Transfer Agreement (MTA). All sequence data will be deposited under NCBI Bioproject Accession PRJNA343262.

### Funding

This project was supported by National Research Initiative grants number 200835205-04687 and 2008-35205-18864 from the USDA Cooperative State Research, Education and Extension Service and National Research Initiative grants number 2009-65205-05635 and 2013-68004-20364 from the USDA National Institute of Food and Agriculture.

### Authors’ contributions

JLH JED and RDS JFT designed and conducted the study. JFT and RDS collected samples and processed the genotypes. RDS processed NGS data. JLH conducted bioinformatic analyses of genotypes and sequence variants. JLH and JFT wrote the manuscript. JFT RDS and JED edited the manuscript.

### Competing Interests

Authors have no competing interests to declare.

## Acknowledgements

We gratefully acknowledge the provision of semen samples from Angus breeders and semen distributors. We appreciate the financial support of the American Angus Association, the Australian Angus Association, The New Zealand Angus Association and the Argentine Angus Association to sequence the genomes of bulls used in this study.

